# Lateral Prefrontal Theta Oscillations Causally Drive a Computational Mechanism Underlying Conflict Expectation and Adaptation

**DOI:** 10.1101/2024.04.30.591918

**Authors:** María Paz Martínez-Molina, Gabriela Valdebenito-Oyarzo, Patricia Soto-Icaza, Francisco Zamorano, Alejandra Figueroa-Vargas, Patricio Carvajal-Paredes, Ximena Stecher, César Salinas, Antonie Valero-Cabré, Rafael Polania, Pablo Billeke

## Abstract

Adapting our behavior to environmental demands relies on our capacity to perceive and manage potential conflicts within our surroundings. While evidence implicates the involvement of the lateral prefrontal cortex and theta oscillations in detecting conflict stimuli, their roles in conflict expectation remain elusive. Consequently, the exact computations and neural mechanisms underlying these cognitive processes still need to be determined. To address this gap, we employed an integrative approach involving cognitive computational modeling, fMRI, TMS, and EEG. Our results revealed a computational process underlying conflict expectation, which correlated with activity in the superior frontal gyrus (SFG). Furthermore, rhythmic TMS in the theta range applied over the SFG, but not over the inferior frontal junction, induced endogenous theta activity, enhancing computations associated with conflict expectation. These findings provide compelling evidence for the causal involvement of SFG theta activity in learning and allocating cognitive resources to address forthcoming conflict stimuli.

**Significant Statement:** Alterations in the processing of expectations of conflict events have been associated with several neuropsychiatric disorders that significantly affect the quality of life for many individuals. This article describes a cognitive computation underlying the conflict expectation and its causal neural mechanism involving theta brain activity in the superior frontal gyrus (SFG). Thus, unraveling this mechanism holds promise for developing interventions to address cognitive alterations related to anticipation of conflict events in neuropsychiatric disorders, improving overall cognitive function and quality of life.

## Introduction

In our daily lives, we often encounter environmental challenges that require us to navigate and anticipate conflict situations—for example, determining when to adjust our driving speed to safely maneuver through traffic signals indicating potential hazards, even when running late for work. In such situations, it is necessary to learn about the possibility of difficult situations and allocate cognitive resources to deal with them.^1–3^ Crucially, we can integrate previously occurring events to anticipate the need for behavioral control; this process is commonly called conflict adaptation or proactive cognitive control.^4,5^ Conflict adaptation involves integrating past events and experiences to adjust goal-directed actions and improve the control required for upcoming stimuli. Clinical literature has identified that alteration in this processing is a component of multiple pathways that contribute to the development of symptoms in various neuropsychiatric disorders^5,6^ such as attention deficit hyperactivity disorder,^7^ depression,^8^ bipolar depression,^9^ obsessive-compulsive disorder,^10^ Parkinson’s disease^11^, and schizophrenia.^12–14^ However, the conventional evaluation of this process, using, for example, the sequence effects in cognitive tasks, does not show robust evidence, presenting disparate results across multiple experimental studies.^15–18^ Additionally, simple concepts such as proactive cognitive control may oversimplify a more intricate and integrative processing required to anticipate and navigate environmental stimuli.^19–22^ Hence, while conflict adaptation is widely acknowledged as crucial for our adaptation to real-life situations, the evidence remains contradictory, primarily due to a lack of comprehensive computational understanding of how this processing occurs.^23–25^ Hence, unraveling the underlying brain mechanisms is critical to rationally developing intervention strategies. Despite ongoing research efforts, the exact computations and neurobiological mechanisms that underlie the expectation of conflict are still not fully understood.

Neuroimaging studies have underscored the significance of the lateral prefrontal cortex in situations where organisms require flexible cognitive control.^26^ Lateral prefrontal activity correlates with behavior regulation according to the current perceptual context and the temporal sequences in which stimuli occur.^2,27^ Context adaptation and its deficit have been associated with the dorsolateral prefrontal cortex.^14,28^ Neurophysiological studies have remarked on the role of oscillatory activity in theta range in several aspects of cognitive control, including its deficit in pathologies.^7,29–34^ Oscillatory activity in the theta band over the frontal midline is associated with detecting, communicating, and implementing cognitive control.^29,35^ Theta-band oscillations have been tied to adaptive control mechanisms during response conflict.^35^ In addition, conflict anticipation positively correlates with the low-theta band’s power.^36^ Despite this evidence, the specific computational processes that underlie conflict anticipations and adaptation and whether they are causally implemented by theta activity in prefrontal areas remain unclear.

Here, we tested the hypothesis that theta oscillatory activity in the lateral prefrontal cortex plays a causal role in computing conflict expectation and adaptation. We anticipate that at the behavioral level, reaction time will increase as the expectation of a conflicting event rises in accordance with specific learning computations. The conflict expectation should be reflected in an adaptation of cognitive control during conflict stimuli. At the neural level, we anticipate that the lateral prefrontal brain network correlates with the conflict expectation computations. Additionally, we hypothesized that non-invasive brain stimulation at theta frequency over the lateral prefrontal cortex enhances the effect of conflict expectation. We conducted three sequential experiments using tasks that induce conflict expectations to test these hypotheses. First, we modeled reaction times to identify a candidate computation process underlying conflict expectations. Second, we used fMRI to identify brain areas correlating with this candidate computation. Finally, in a TMS-EEG experiment, we tested the causal role of theta activity in the identified brain areas in solving conflict expectations (see Supplementary Fig. 1).

## Results

### Modeling Conflict Expectation

An illustrative example of situations that generate conflict expectation is provided by the Go-Nogo (GNG) task (Fig. 1B). This task requires inhibitory responses to rapid and infrequent Nogo stimuli. In such a context, as more consecutive Go stimuli occur, there is a higher probability that a Nogo stimulus will occur in the next trial. The expectation of the occurrence of conflict stimuli involves controlling the response, generating longer reaction times for Go stimuli. Such expectations can be based on specific context features, such as the number of preceding Go stimuli.^7,37^ Thus, the context of recent stimuli can be integrated to improve the inhibitory response to Nogo stimuli.^36^ We anticipate that subjects will adapt their cognitive control based on the estimated probability of encountering conflicting stimuli (Nogo stimuli) and their expectation in the context of previous stimuli. To investigate this, and building on prior research,^7,37^ we tested several models using a sample of 30 subjects who performed the task.

**Fig. 1:**
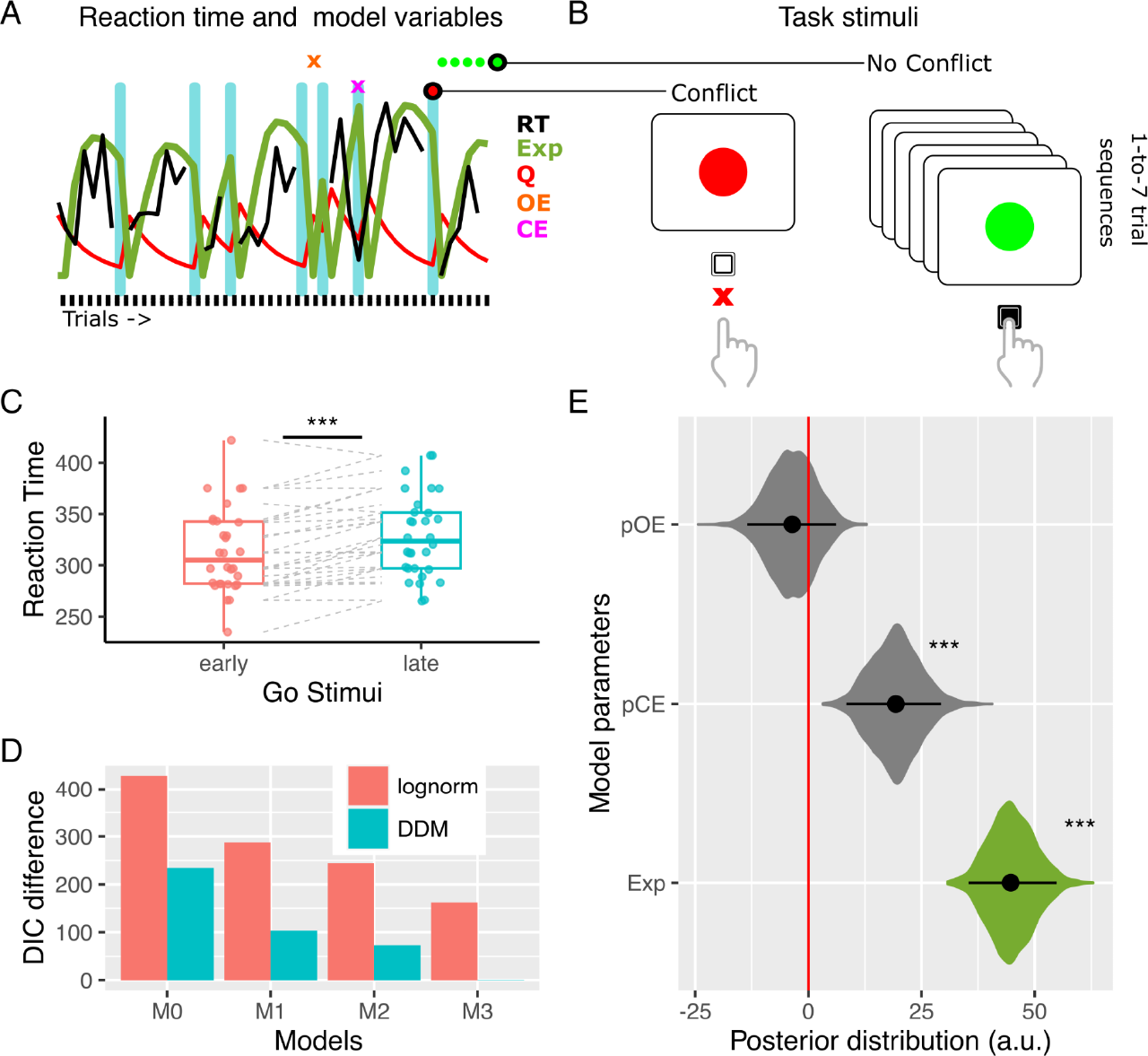
Behavioral analysis of GNG Task. A) Single-trial example of reaction time and model variables. B) GNG Task. C) Reaction time comparison between the first Go trials (early) and the last Go trial (late) of a sequence. D) Models comparison. E) Posterior distribution of model parameters. CE: Commission Error. DIC: Deviance Information Criteria. Exp: Conflict Expectation. OE: Omission Error. pCE: Previous Commission Error. pOE: Previous Omission Error. RT: Reaction Time. See also Supplementary Table 1.

We used reaction time as a proxy for cognitive control and assessed two linking functions for each model: a lognormal and a Wiener (Drift Diffusion Model^38^) distribution. In a null model (M0), we included all relevant regressors except conflict expectation. These regressors encompassed the anticipated slowing of response following errors, categorized as previous commission errors (pCE) and previous omission errors (pOE).

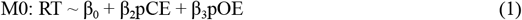

Then, in the first cognitive model (M1), we hypothesize that the expectation of conflict stimuli (Exp) linearly increases with the number of consecutive Go stimuli in a sequence (nSeq) of stimuli, such that Exp = nSeq.

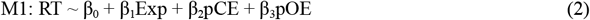

In the second model (M2), the expectation of conflict stimuli (Exp) is determined by the rational calculation of the probability of the occurrence of the given sequence of Go stimuli, as expressed in the following equation.

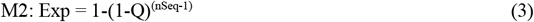

In the preceding equation, Q represents the rate of occurrence of conflicting stimuli, which, for the M2 model, was set to 0.25 (the programmed rate of Nogo stimuli). The −1 exponent indicates the impossibility, known by the subject, of the occurrence of two consecutive Nogo stimuli. In other words, the probability of Nogo stimuli occurring in the first position in the sequence is zero.

In the third model (M3), we expected that participants learn the rate of occurrence of conflict stimuli, Q, using a learning algorithm as follows.

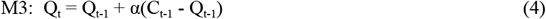

In the preceding equation, α represents the learning rate, and C_t-1_ indicates the presence (C_t-1_ = 1) or absence (C_t-1_ = 0) of a conflicting stimulus in the prior trial.

### Behavioral Experiment: Conflict Expectation Effect

Based on the preceding rationale, we initially analyzed the impact of expectation modulation on behavior in a sample of 30 healthy participants. To accomplish this, we compared the RT for the first two “go” stimuli in a sequence with the subsequent three “go” stimuli (i.e., 1st and 2nd vs. 3rd, 4th, and 5th stimuli). This division ensured a relatively balanced distribution of trials for comparison (52.3% vs. 47.7%). Our results revealed that the RTs for the initial trials were significantly faster and less accurate than the final trials (RT: mean differences −14.3 ms, confidence interval (CI) = [-19.9 −8.6], Wilcoxon test, p = 8.8e-5, effect size r = 0.7; accuracy rates: differences 0.08, CI = [0.06 0.1], Wilcoxon test, p = 1.8e-6, effect size r = 0.8). These results could indicate that the subjects adjusted their cognitive control, altering the speed-accuracy trade-off.

We then applied the models described in the previous section to evaluate which formulation best explained the mechanism underlying the increase in RT associated with the expectation of conflict stimuli (Table 1). Among the different models tested, the best-fitted model was M3 with the drift-diffusion model (DDM, e.i., Wiener distribution, see Table 1 and Supplementary Table 1). In this model, the regressor of conflict expectation had a significant impact on reaction time (β_1_ Expectation mean = 44.7, 95% high-density interval (HDI) of [35 54], p_MCMC_ <0.001, p_MCMC_ is a p-value derived by comparing the posterior distributions of the estimated parameters sampled via Markov Chain Monte Carlo, see Methods). Additionally, commission error, which refers to the previous error in a “Nogo” stimulus, was found to significantly increase the reaction time for the “Go” stimuli (β_2_ pCE mean = 19.3, HDI = [8 29.3], p_MCMC_<0.001). However, no significant effects were found for a previous omission error (beta pOE mean = −3.6, HDI = [-13 6], p_MCMC_=0.49, Fig. 1D). The posterior distribution of the learning rate was a mean of 0.29 and an HDI of [0.22 0.37]. Then, we test if the expectancy of conflict stimuli would significantly influence inhibitory control accuracy, as measured by the proportion of successful Nogo inhibition. Results showed a significant positive effect of expectation on the subsequent inhibitory control accuracy (β_a1_ mean = 1.1, HDI = [0.17 1.98], p_MCMC_=0.012).

**Table 1:** Summary of the models and their indicators of adjustment.

| **Model** | **Linking function** | **Free parameters** | **DIC** | **LOO** |
| --- | --- | --- | --- | --- |
| M0<br>null | lognormal | 4 (3 betas, sigma) | 427.1 | 480.1 |
|  | Wiener | 5 (3 betas, drift, tau) | 233.0 | 257.5 |
| M1<br>linear | lognormal | 5 (4 betas, sigma) | 286.7 | 326.4 |
|  | Wiener | 6 (4 betas, drift, tau) | 102.1 | 109.6 |
| M2<br>Exp | lognormal | 5 (4 betas, sigma) | 243.6 | 279.5 |
|  | Wiener | 6 (4 betas, drift, tau) | 71.7 | 71.3 |
| M3<br>Exp+LR | lognormal | 6 (4 betas, alpha, sigma) | 161.5 | 24.1 |
|  | Wiener | 7 (4 betas, alpha, drift, tau) | 0 (reference) | 0 (reference) |

In summary, individuals adjust their RT according to the expectancy of the conflict stimuli—such processing, which could reflect cognitive control allocations, involved learning from past experiences in the environment.

### GNG Behavioral replication sample

We carried out a replication of the behavioral analysis in an independent sample of 57 participants. The task presented two modifications to control possible confounding factors. First, the sequences were not restricted to a maximum of seven ‘Go’ stimuli, and the sequence formation was completely randomized following a 0.25 probability of ‘Nogo’ occurrence (see Methods). Secondly, the stimuli were letters (O and X) rather than colors to avoid possible color bias, and they were counterbalanced to serve as Go or No-Go stimuli in the two blocks for each subject. We replicated all the findings of the original sample (Supplementary Fig. 2). The early ‘Go’ stimuli were faster (mean difference = −21 ms, CI = [−26, −15], Wilcoxon test, p = 7e-11, effect size r = 0.78), and the ‘Nogo’ stimuli preceded by short ‘Go’ sequences were less accurately inhibited (mean rate difference = 0.09, CI = [0.07, 0.11], Wilcoxon test, p = 1e-10, effect size r = 0.86). The M3 model with DDM fitted the data best, and the Exp regressor significantly differentiated from zero (β_1_ Expectation mean = 51.7, 95% HDI: [51 62], p_MCMC_ <0.001, see Supplementary Fig. 2).

### MRI Experiment: Modeling MSIT

Our next objective was to identify the brain areas responsible for computing the expectation of conflict stimuli. We utilized the fMRI technique due to its high spatial resolution to achieve this goal. However, the slow dynamics of hemodynamic responses precluded using fast stimuli presentation tasks and, therefore, the generation of sequence effects, as seen in the GNG task.^37^ To overcome this limitation, we designed a task and analysis based on the same rationale as the behavioral experiments adapted for fMRI. Specifically, we used the MSIT task, which involved variable sequences of congruent and incongruent stimuli, to obtain a balanced number of trials for the contrast (see Methods and Results sections below). Furthermore, for the analysis, we used conflict expectation as equivalent to the probability of conflict (i.e., Exp = Q, instead of Eq. 3, see Fig. 2A) as has been done in other research studies with large interstimulus intervals.^36,39^ Following this rationale, we tested whether subjects adjusted their reaction time according to the expectation of the occurrence of conflict stimuli. First, as in the behavioral experiment, we separated the first two stimuli in a sequence with the subsequent three stimuli (i.e., 1st and 2nd vs. 3rd, 4th, and 5th stimuli) in a sample of 26 participants. Following the behavioral model utilized in the GNG task, we expect that in the early trial for congruent sequences, subjects should expect more conflict than in the late trials, and vice versa for incongruent sequences. Accordingly, the trials with more predicted conflict expectation present slower RT (mean differences: 70.6 ms, CI = [85.9 55.3], p=2.9e-8, effect size r=0.87, Fig. 2C). Accuracy modulation was found only for incongruent sequences representing conflict stimuli (mean rate differences: 0.06, CI = [0.03 0.1], p=0.0007, effect size r=0.59). We then applied the M3 model using the drift-diffusion model. The results showed that the probability of conflict significantly increases the RT in the MSIT task (β_1_ Expectation mean = 1.07, HDI: [0.74 1.43], p_MCMC_<0.001, Fig. 2C). As expected, conflict stimuli are slower than no conflict stimuli (β Conflict mean = 0.32, HDI: [0.22 0.41], p_MCMC_<0.001). To summarize, in the MSIT task, it is possible to identify homologous expectation processing with a similar computational mechanism to that specified in the GNG task.

**Fig. 2:**
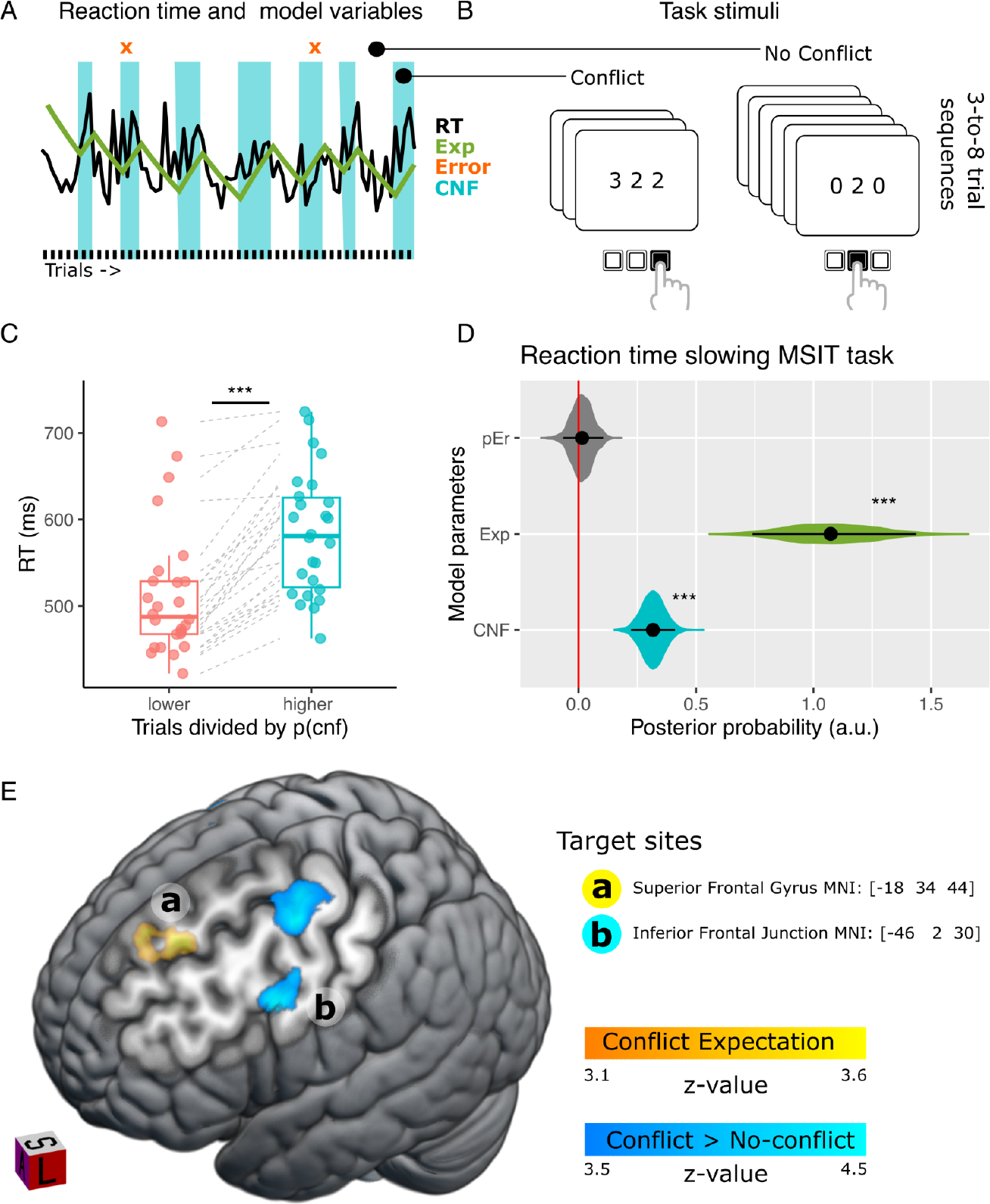
MRI Experiment Results. **A**. Example of trial-by-trial variation in the reaction time and model variables. **B**. MSIT Task stimuli. **C**. Comparison of reaction times between trials with predicted lower and higher conflict expectations per sequence. Each point represents the mean of a subject. **D**. Posterior distribution of model parameters. **E**. Brain activity is associated with conflict expectation in the lateral prefrontal cortex (yellow) and conflict stimulus processing (blue). Lowercase letters “a” and “b” indicate the target sites for TMS experiments. pEr: Error in the prior trial, Exp: Conflict Expectation, CNF: Conflict trial, p(cnf): Predicted probability of conflict, RT: Reaction Time.

### MSIT Behavioral replication sample

We carried out a replication of the behavioral analysis in an independent sample of 63 subjects. We replicated all the findings of the original sample (Supplementary Fig. 3). The trials with predicted less conflict expectation within a sequence were faster for both sequences (mean difference = −58 ms, CI = [−69 −46], Wilcoxon test, p = 6e-12, effect size r = 0.86), and less accurate for incongruent sequences only (mean rate difference = 0.02, CI = [0.007 0.04], Wilcoxon test, p = 0.01, effect size r = 0.3). The Exp regressor significantly differentiated from zero (posterior distribution, β_1_ Expectation mean = 1.02, HDI: [0.68 1.37], p_MCMC_ < 0.001, Supplementary Fig. 3).

### MRI Experiment: Brain activity correlating with expectancy of conflict

To evaluate the brain areas in which activity correlated with the expectation of conflicting stimuli, we modeled the BOLD signal using the conflict expectation for each trial and the presentation of a conflicting or non-conflicting stimulus in the current trial. This analysis showed that the regressor of expectation (Exp) exhibited a significant correlation with bilateral activity in the SFG (see Fig. 2C) and the medial parietal cortex (see Supplementary Table 2). Furthermore, the contrast between conflict and non-conflict stimuli presented a significant correlation with the activity in the frontal eye field (FEF) and the inferior frontal junction (IFJ) in the lateral prefrontal cortex in both hemispheres, as well as other brain areas (see Fig. 2C and Supplementary Table 2).

### EEG-TMS Experiment

#### SFG TMS at theta frequency enhances conflict expectancy during the GNG Task

Prior work has demonstrated that theta oscillation and lateral prefrontal cortex activity correlate with proactive cognitive control.^7,14,40^ Hence, based on our fMRI results, we aimed to investigate the causal relationship between theta activity in the SFG and the conflict expectation. To this end, we conducted a TMS-EEG experiment using TMS stimulation before each 20-stimuli run of the GNG task (Fig. 3A). We selected a 5Hz frequency for stimulation and used arrhythmic TMS and sham stimulation as control. Considering our fMRI results, we stimulated two different areas, the left SFG (MNI: [-18 34 44], peak coordinate for the Expectation regressor, see Fig. 2) and the left IFJ (MNI: [-46 2 30], peak coordinate for the Conflict regressor, see Fig. 2), in separate sessions to assess their respective involvement. We targeted the SFG to investigate its role in the conflict expectation while controlling for potential effects on inhibitory control by stimulating the IFJ.^41,42^ We chose the left hemisphere because the peaks of BOLD activity were in this hemisphere. Additionally, previous research has indicated that clinical samples demonstrating alterations in proactive cognitive control often exhibit reduced left-lateralized target activity.^7^

**Fig. 3.**
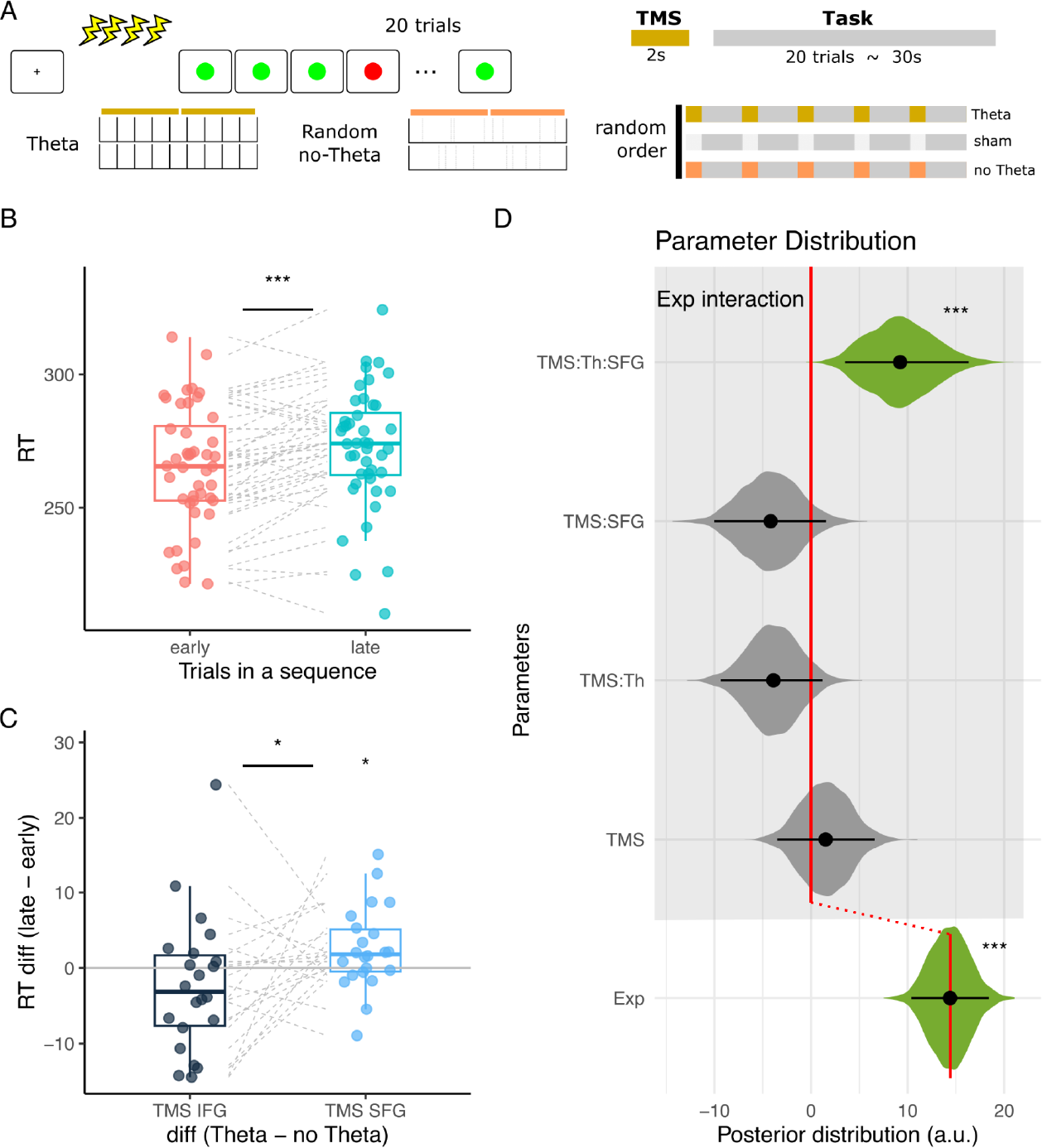
TMS experiment behavioral results. **A**. Experimental design of the GoNogo Task during EEG-TMS sessions. **B**. Reaction time (RT) modulation by the trial position in a sequence. **C**. The modulation of the effect of trial position by the TMS stimulation. **D**. Posterior distribution of model parameters. Black dots represent the mean of the distribution, and black lines represent the 95% high-density intervals. The colored areas represent the complete posterior distribution. * indicates p<0.05, ** p<0.01,***p<0.001. RT: Reaction Time. TMS: Transcranial Magnetic Stimulation. Th: Theta Rhythm. SFG: Superior Frontal Gyrus.

Using the peak activity in the SFG as our target site, we explored the causal relationship between theta activity and the conflict expectation. Our behavioral results revealed that participants exhibited slower RT as the conflict expectation increased. The first trials of a sequence (the 1st and 2nd stimuli) were faster than late trials (the 3rd, 4th, and 5th stimuli; mean difference −7.1 ms, CI = [-10.2 −4]; Wilcoxon test p = 0.0001, n = 44, r=0.57, Fig. 3B). This effect was significantly greater as the theta TMS stimulation was applied in the SFG in comparison with arrhythmic TMS stimulation (mean difference = −2.5 ms, CI = [-4.9 −0.08], Wilcoxon test p = 0.04, r=.43, n=22, Fig. 3C). No effect was found when theta TMS stimulation was applied in the IFJ (mean differences: 2.2 ms, CI=[-1.7 6], Wilcoxon test p = 0.2, r=0.2, n=22, Fig. 3C). Thus, the theta effect was significantly greater for the SFG than for the IFJ TMS stimulation (mean differences = −4.8 ms, CI=[−9.4 −0.2], Wilcoxon test p = 0.04, r=.44, n=22, Fig. 3C).

We then tested the cognitive model M3, showing the same effect. Specifically, we found that participants during sham stimulation in both sessions showed similar behavior to those in the behavioral experiment, increasing their reaction time in response to the conflict expectation (β_1_ Exp mean: 14.3, HDI: [10.3, 18.3], p_MCMC_ < 0.001). Regarding TMS stimulation, we observed that the main effect of TMS, regardless of its frequency or target site, did not significantly impact participants’ RT (β_8_ TMS mean: 0.4, HDI: [-2.4, 3.0], p_MCMC_ = 0.76; β_4_ Exp*TMS mean: 1.52, HDI: [-3.5 6.5], p_MCMC_ = 0.56). However, we found that theta stimulation over the SFG specifically resulted in an expectation-related increase in RT. This was evidenced by a significant interaction between expectation, TMS, theta, and the SFG (β_7_ Exp*TMS_theta*SFG_ mean= 9.3, HDI= [3.5 16.3], p_MCMC_ = 0.0006; see Supplementary Table 4 and Fig. 3B). Concerning the accuracy, the TMS stimulation had no direct effect on the accuracy (β_11a_ TMS_SFG_ mean = 0.042, HDI = [-0.25 0.34], p_MCMC_=0.7). However, we did observe that the impact of expectation given by TMS_theta*SFG_ stimulation influences the accuracy of following Nogo stimuli (β_a7_ Exp*TMS_theta*SFG_ mean= 0.27, HDI = [0.05 0.49], p_MCMC_=0.002; β_1a_ Exp: mean= 0.44 HDI = [0.19 0.68], p_MCMC_<0.001). The findings suggest that theta stimulation over SFG may precisely modulate cognitive processes related to conflict expectation.

#### TMS stimulation in the SFG increases endogenous theta oscillation

We conducted a power analysis of the EEG signal to assess whether TMS theta stimulation induces oscillatory brain activity. We examined the time windows following TMS stimulation and just before task stimulus presentation, revealing a specific effect of TMS stimulation. Rhythmic theta stimulation over the SFG significantly enhanced theta activity compared to sham stimulation (Fig. 4A). We found the same tendency compared to arrhythmic stimulation, but the effect did not survive cluster correction (Fig. 4B). We also observed higher theta activity than theta stimulation over the IFJ (Fig. 4C). Source analysis indicated the involvement of the right lateral prefrontal cortex. The gray rectangle in Fig. 4 represents the common area revealed by conjunction analyses, as observed in the contrasts Theta TMS > sham in SFG and Theta TMS in SFG > Theta TMS in IFJ. Notably, a shared modulation was identified in a cortical region where we had observed modulation in the fMRI experiment, representing the contralateral stimulation area. Overall, TMS theta stimulation can generate brain oscillatory activity in the theta range contralateral to the stimulated brain region in the SFG.

**Fig. 4:**
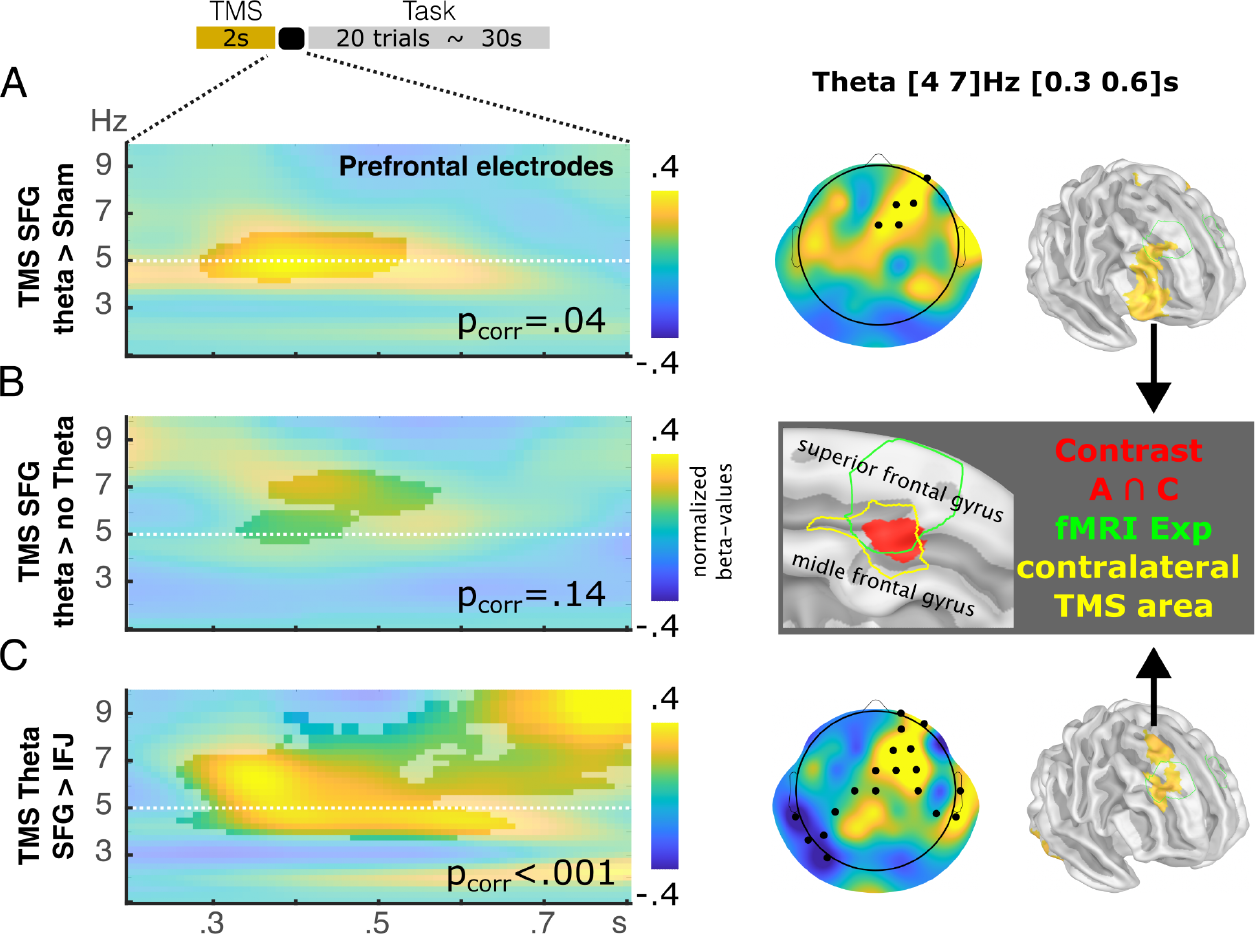
Oscillatory effects post-TMS. **A**. Power increases after theta TMS stimulation over the SFG compared to sham stimulation. The left panel shows the time-frequency chart for a second after the last TMS pulse and before the first Go stimuli. The right panel shows the scalp distribution and source estimation of theta modulation. **B**. Power increases after theta TMS stimulation over the SFG compared to no theta stimulation. No modulation in source space survived multiple comparison corrections. **C**. Power increases after theta TMS stimulation over the SGF compared to theta TMS stimulation over the IFJ. The scalp distribution and source estimation reveal the main modulation of the left lateral frontal cortex. The gray rectangle shows the conjunction analysis for the contrasts shown in A and B. The green line represents the BOLD modulation for the Exp regressor in fMRI experiments (see Fig. 2). The yellow line represents the homologous contralateral area of stimulation (Braintomme atlas area: A9/46d^43^). TMS: Transcranial Magnetic Stimulation. SFG: Superior Frontal Gyrus. IFJ: Inferior Frontal Junction.

#### The conflict expectation modulates the amplitude and the phase-amplitude coupling of the lateral frontal oscillatory activity

We next analyzed the brain activity during task execution. We conducted a power analysis of the electrophysiological recordings and correlated it with the conflict expectation predicted by the behavioral model. These analyses indicated a statistically significant modulation in theta activity associated with the conflict expectation. During the ‘sham’ conditions, a post-stimulus modulation in theta oscillations related to conflict expectation was observed at the frontal electrodes in response to the ‘go’ stimulus (see Fig. 5A). Source analysis revealed that this modulation was localized in the right lateral prefrontal region. Previous evidence suggests prefrontal areas exert control functions through phase-amplitude coupling (PAC)^44^. For instance, studies have demonstrated that the low-frequency phase in the delta/theta range modulates the amplitude of alpha/beta oscillations, reflecting control signals over other cortical regions, such as motor and premotor areas.^45–47^ Hence, we assessed PAC modulation that could reflect increased control associated with heightened expectation of conflict. We calculated the Envelope-to-Signal correlation in a time window following stimulus presentation and correlated this with the expectative of conflict stimuli. This analysis revealed that the expectative of conflict correlated with an increased PAC between delta/theta and alpha/beta oscillations (Fig. 5B). This modulation was placed over the right central electrodes that were compatible with the motor and premotor areas.

**Fig. 5:**
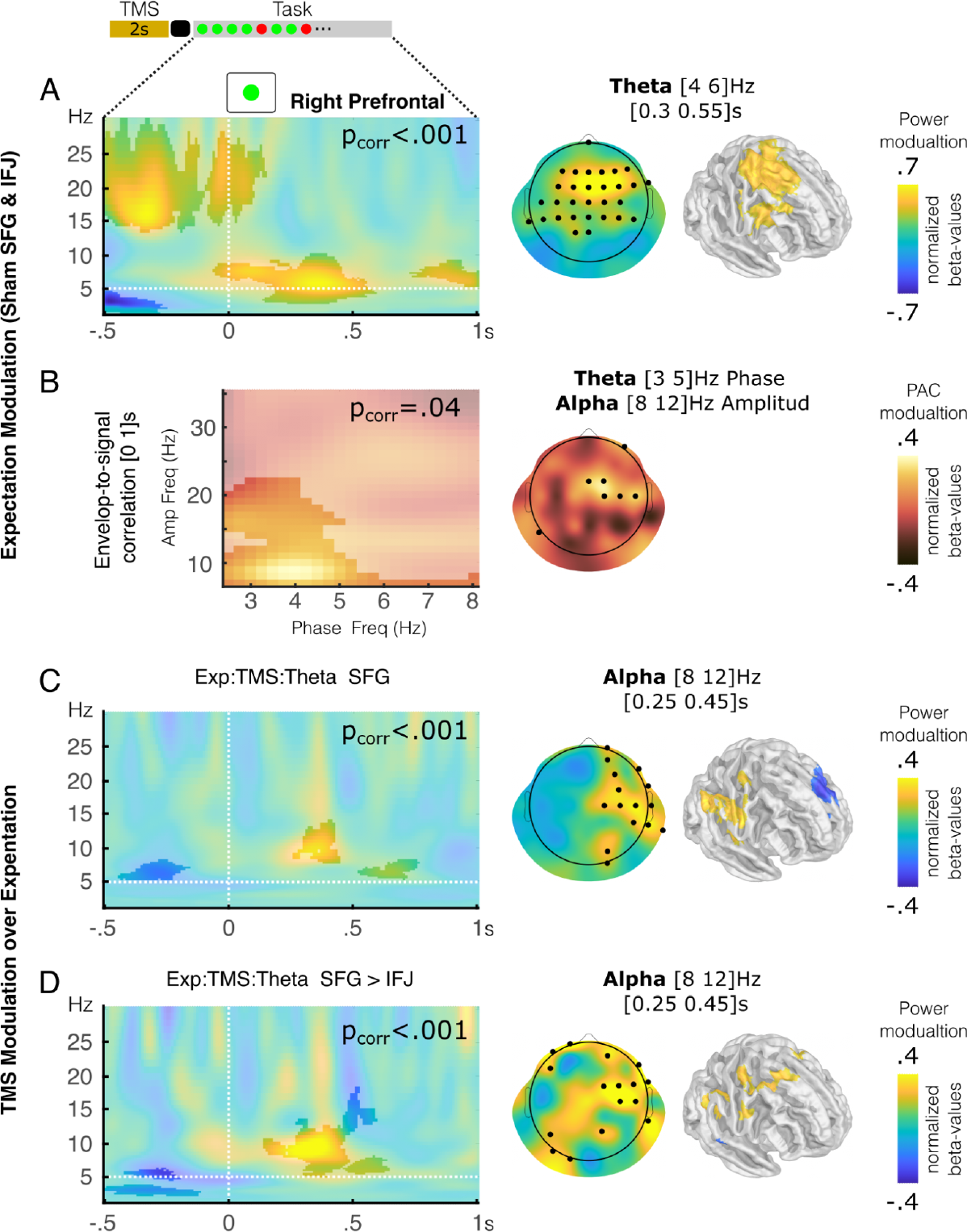
Oscillatory modulation related to conflict expectation and TMS effects. **A**. Oscillatory activity associated with the expectation of conflict in sham conditions is depicted. The left panel illustrates the time-frequency chart over frontal electrodes, while the right panel displays the topographic and cortical surface distribution of theta modulation. **B**. Phase amplitude coupling during the second following Go stimuli modulated by the expectation of control. **C**. Oscillatory activity related to the conflict expectation and TMS theta interaction during SFG stimulation. **D**. Oscillatory activity related to the conflict expectation and TMS theta interaction for the contrast between the SFG and IFJ stimulation. **A-D**. The highlighted areas represent significant modulation. The electrodes highlighted indicate those with significant modulation. The vertical dotted lines denote the appearance of Go stimuli, and the horizontal dotted line represents the frequency of theta stimulation. SFG: superior frontal gyrus, IFJ: inferior frontal junction.

#### TMS stimulation modulates theta oscillations in lateral prefrontal regions related to the conflict expectation

Next, we explore TMS’s modulation in electrophysiological activity related to conflict expectations. Exploring the interaction among Expectation, TMS, and theta stimulation, we found a positive modulation in frontolateral electrodes for both theta and alpha oscillations generated by TMS over SFG (Fig. 5C). Contrasting the preceding interaction between the SFG and IFJ TMS interaction, a similar modulation was detected involving theta and alpha oscillations (Fig. 5D). Scalp topographies and source analysis of this modulation reveal the prominent participation of alpha modulation in the right frontolateral region involving premotor and motor areas and the parietal region. No modulation was found to TMS in the correlation between PAC modulation and conflict expectations.

## Discussion

Improving our understanding of anticipation and adaptation to conflict stimuli is pivotal for implementing therapies targeting such processing across diverse neuropsychiatric disorders. Employing an integrative methodology that combines computational modeling, neuroimaging, transcranial magnetic stimulation, and electroencephalography, our goal was to uncover the mechanisms underlying these cognitive processes. Our results described computational and neurobiological mechanisms underlying the formation of expectations related to conflicting events, as reflected in reaction times, and involving theta activity in the lateral prefrontal cortex in a causal manner.

Our behavioral modeling revealed that participants adapt their reaction times in anticipation of conflict stimuli, as evidenced by slower responses following a higher expectation of such events. This adaptive behavior suggests a flexible mechanism wherein individuals dynamically allocate cognitive resources based on learned expectations.^48,49^ Thus, participants anticipate conflict, suggesting that cognitive processes associated with conflict monitoring and resolution were engaged before the conflict stimulus presentation. Recent research indicates that the dynamic process of learning from conflicting stimuli in the environment is correlated to multidimensional representations within neuronal ensembles.^39^ Furthermore, indicators of conflict occurrence may be associated with accumulating learning from conflicts over time, influencing future responses to similar conflicting stimuli.^39,50^ Interestingly, this learning has specific task-dependent and general doming mechanisms.^39^ Our cognitive model posits analogous patterns, indicating the existence of a domain-general computation that remains consistent across tasks and adapts to learn from conflicting environmental stimuli. Conversely, specific expectations, like the sequence effect, undergo adaptation tailored to the particular demands of the task, such as adjusting to interstimulus intervals.^37^ Hence, the cognitive computation model proposed here can be understood as a general domain processing that could occur in different situations and be utilized in various ways depending on the specific context and demands. Thus, this computational process can be tested in different contexts and has the potential to reconcile contradictory evidence and theoretical propositions to understand conflict adaptation and flexible cognitive control^23,24^, especially concerning understanding various neuropsychiatric pathologies.^51^

Our fMRI results further elucidated the relationship between conflict expectation and frontal activity. The observed engagement of distinct regions within the lateral prefrontal cortex in response to conflict stimuli and conflict probability is consistent with prior neuroimaging research, which has consistently shown the frontal brain areas’ involvement in conflict resolution, conflict monitoring, cognitive control, and response inhibition.^2,27,52,53^ In the study of perceptual decision-making, a dissociation between various regions of the lateral prefrontal cortex has been linked to distinct cognitive computations^54,55^. Our results highlight the differential causal role of the SFG and IFJ in mediating the impact of conflict expectation on behavioral performance. The SFG is a region implicated in different tasks, including integrating working memory and inhibitory control to solve cognitive tasks^56^ and accumulating and valuing sensory evidence from the environment^55^. Moreover, alteration in SFG activity is associated with disorders such as bipolar depression^9^ and schizophrenia.^12,13^ On the other hand, the IFJ is essential for executive functions^57^ like control of action, attention, and memory,^58^ and its bilateral lesion causes the dysexecutive syndrome.^57^ Interestingly, the interaction between these two areas is related to the flexible use of proactive or reactive strategy.^59^ Thus, the lateral frontal areas and their associated networks play a crucial role in abstracting diverse environmental features to estimate the probability of encountering difficult situations. This process involves computations akin to learning algorithms that dynamically adjust real-time control mechanisms to adapt behavior.

Our EEG-TMS results shed light on the neurobiological implementation of the described computational mechanism. TMS stimulation targeting the left SFG at theta frequency enhances conflict expectation during the GNG task. Using repetitive TMS to increase cortical excitability, a study showed that stimulation of the left lateral prefrontal cortex, rather than the right, enhances proactive cognitive control.^60^ In addition, evidence has shown that using TMS with specific experimental designs has improved other functions such as verbal memory^61^ and working memory.^62,63^ Although TMS over the left frontal cortex has been proposed to enhance frontal functions such as flexibility and conflict adaptation, contradictory results exist.^64^ The variability in outcomes observed in studies concerning the stimulation of the left lateral prefrontal cortex may stem from a deficiency in precise computational mechanisms governing cognitive processes and the inability to target specific neuronal activities accurately. Our experimental approach enables us to intervene with precise neurobiological activities. Previous research has shown that TMS can induce theta activity during task execution.^62^ In this line, we observed an increase in theta activity following TMS stimuli, demonstrating that exogenous stimulation with TMS can interfere with and augment endogenous oscillatory activity, as observed in other studies.^65^ In our work, TMS effectively produces an oscillatory intervention in the prefrontal network.

The source analysis of the oscillation induced by TMS shows contralateral activity. This activity could be explained by the fact that TMS does not have a local effect but a network effect.^66,67^ This non-local effect depends on structural rather than functional connectivity.^68^ Stimulation with TMS in one area generated secondary stimulation in its connected areas, demonstrating an effect in its homologous contralateral area, as seen in studies with TMS in parietal regions,^69^ in the auditory cortex,^70^ and in the motor cortex.^71^ TMS depolarizes cortical neurons, and the activity expands to all connected areas. This effect of TMS would be the basis for rehabilitation therapies,^72^ being able to make interventions at the cortical level, producing an effect in deeper areas.^73^ In this context, it is relevant to consider individual differences in functional brain connectivity, which could interfere with the effects of TMS.^74^ Additionally, at the site of direct stimulation, the depolarization of different types of neurons in the local circuit generates a non-physiological functioning that may explain unclear effects on activity observed in some research at the target site.^75^ However, in the connected area, the stimulation induced by the depolarization of pyramidal neurons at the target site can yield a more physiological estimation of the output circuit, which can explain the activity and entrainment of oscillatory theta activity found in the contralateral area in our experiment.

Frontal theta activity has been related to diverse cognitive functions, including conflict evaluation and monitoring. Our results indicated that the conflict expectation modulates the power of theta activity in the right lateral prefrontal cortex according to the cognitive demands of the task. As a complex mechanism, cognitive control involves theta oscillations and the increase of theta gamma phase-amplitude coupling (PAC) in the dorsomedial and dorsolateral prefrontal cortex^76^ and the orbitofrontal cortex.^77^ Hence, our cross-frequency analysis showed that conflict expectation also correlated with an increased PAC between the phase of delta/theta and the amplitude of alpha/beta oscillations. The later modulation could be the mechanism by which cognitive control is exerted towards the motor and premotor cortex,^45^ relating to inhibitory control.^47^ Additionally, delta-beta coupling has been proposed as a marker for anticipatory anxiety in pathologies such as social anxiety disorder.^78^ Interestingly, TMS modulates theta and alpha activity in the lateral prefrontal cortex during task execution, increasing conflict expectations’ behavioral effect. The results show that these oscillations are a key component of algorithmic processing to face conflict tasks, as has been demonstrated and postulated in other cognitive processes such as the attentional network^79^ and working memory.^80^ The contrast between TMS stimulation at theta rhythm in different regions of the lateral prefrontal cortex further demonstrates the specificity of the SFG in this computational process. Hence, considering all EEG results, the theta and alpha oscillations in a prefrontal network could be the mechanism by which the computation of conflict expectation is implemented in the brain.

Alterations in the expectancy of conflict events have been associated with neuropsychiatric disorders that profoundly impact the quality of life for many individuals. Our results describe a cognitive computation underlying conflict expectation and its causal neurobiological mechanism involving theta activity in the SFG. Given the development of diverse techniques for intervening in brain activity in a non-invasive manner in clinical settings^81^ and the known alteration of brain oscillation in diverse disorders,^7,31,40,82,83^ unraveling this mechanism holds promise for developing interventions to address these cognitive alterations in neuropsychiatric disorders, thereby enhancing overall cognitive function and quality of life.

### Methods Participants

One hundred seventy nine healthy subjects participated in this research approved by the Ethics Committee of the Clinica Alemana - Universidad del Desarrollo, Chile. The research consisted of 221 experiential sessions, structured as follows. In the first experiment, 30 subjects (15 women, ages ranging between 18 and 20 years) participated in the behavioral session and 57 in the replication sample (32 woman, ages ranging between 23 and 60 years). In the subsequent experiments, another sample of subjects participated as follows. In the second experiment, 26 subjects (15 women, ages ranging between 18 and 35 years) participated in the fMRI session and 63 in the behavioral replication sample (33 women, ages ranging between 18 and 37 years); in the third experiment, 24 subjects (13 women, ages ranging between 18 and 35 years) participated in the EEG-TMS sessions. A total of 18 subjects participated in experiments 2 and 3. All participants gave informed consent. Experiments were conducted in the Social Neuroscience and Neuromodulation Laboratory at the Centro de Investigación en Complejidad Social (neuroCICS) at the Universidad del Desarrollo and the Unidad de Imágenes Cuantitativas Avanzadas (UNICA) at the Clínica Alemana de Santiago.

### Experimental design

For the behavioral experiment, subjects participated in the GNG task. With these data, different behavioral models were tested to explain conflict expectancy and variations in reaction time. The best-fitted model was used to analyze the data from experiments 2 and 3 (Fig. 1 and Supplementary Fig. 1). For the second experiment, subjects participated in an MRI session and solved the MSIT task. Imaging data were analyzed by integrating regressors from behavioral modeling of the MSIT task to identify target sites for experiment 3. In the third experiment, subjects participated in two sessions of TMS-EEG (Supplementary Fig. 1) while they performed the GNG task. In one session, the stimulation was delivered over the superior frontal gyrus (SFG); in the other, it was in the inferior frontal junction (IFJ). The stimulation site for each session was randomized, and the sessions were balanced.

### Tasks

#### Go-nogo Task

During the behavioral and TMS-EEG experiments, participants completed the Go-nogo (GNG) task, which involved viewing green or red stimuli on a screen. Participants were instructed to press a keypad button for the green stimuli (Go stimulus) but not to press the button for the red stimuli (Nogo stimulus). In the behavioral experiments, participants completed two blocks, each consisting of 150 trials (Fig. 1). In the TMS-EEG sessions (experiment 3), participants completed four blocks containing three sub-blocks of 100 stimuli. The stimuli were separated into five runs of 20 stimuli each. Before each 20-stimuli run, 10 TMS pulse series were applied for 2 seconds in different patterns: rhythmic (5 Hz), arrhythmic, or sham (Fig. 3). The behavioral results of the GNG task were analyzed by fitting different models to estimate the probability of conflict and the expectation of conflict occurrence. Each stimulus was presented over 300 ms, and the interstimulus interval varied between 900 and 1400 ms. The occurrence of Nogo stimuli was set to 0.25, interspersed within sequences ranging from 1 to 7 Go stimuli.

#### Go-Nogo Task for Replication Sample

Following the same rationale as the previous task, for the replication sample, the Go-Nogo task was presented in two blocks of 150 trials where the stimuli were letters (X or O). They were randomly assigned and counterbalanced so that each participant encountered that a letter representing the ‘Go’ stimulus in one block and the ‘Nogo’ stimulus in the other block, and vice versa. Each stimulus was presented over 300 ms, and the interstimulus interval varied between 900 and 1400 ms. The occurrence of ‘Nogo’ stimuli was set to 0.25, and the length of the sequences of ‘Go’ stimuli was not restricted.

#### MSIT Task

During the MRI experiment, participants were presented with a set of stimuli consisting of three numbers and were asked to identify the number that was different from the other two using a keypad. The task included congruent and incongruent trial sequences of variable duration (3 to 8 trials). During congruent trial sequences, the different number matched its position on the keypad and was flanked by non-task-relevant numbers (zeros). In incongruent trial sequences, which were associated with great difficulty, the different numbers and their positions did not match, and the other numbers were task-relevant (Fig. 2). The occurrence of conflict stimuli was 0.5. Each stimulus was presented for 1000 ms, and the interstimulus interval varied between 1500 and 3000 ms.

### Behavioral analysis

Behavioral analysis was conducted using a Hierarchical Bayesian approach, which leverages the aggregated information from the entire population sample to inform and constrain the parameter estimates for each individual. This approach incorporated two levels of random variation: the trial level (i) and the participant level (s). Models were compared using the Deviance Information Criterion (DIC) and the Leave-One-Out Information Criterion (LOOIC). All the models were fitted to the reaction times (RT) and were adjusted using R and JAGS software. Given its proposed influence on the speed-accuracy tradeoff that underlies cognitive control,^84^ we assumed that all regressors would affect the drift-diffusion model’s boundary, and we set the bias parameter to 0.5. Additionally, we examined the effect of cognitive control (CC) on accuracy by constraining the model to include reaction time in the current trial (t) and accuracy (Acu) of the Nogo stimulus in the subsequent trial (where applicable) as follows.

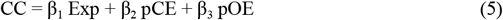

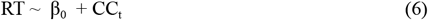

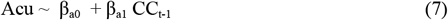

In the TMS-EEG experiment, we used the following model:

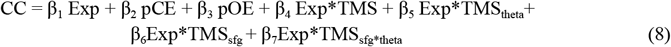

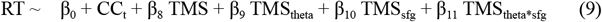

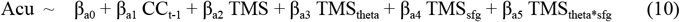

Where TMS represents the main effect of TMS stimulation, TMS_theta_ represents the effect of theta rhythmic stimulation, TMS_sfg_ the stimulation in SF, and TMS_theta*sfg_ represents the specific effect of theta stimulation over SFG.

All β parameters were parameterized using normal distributions, while the α parameter (learning rate, equation 4) was parameterized using a beta distribution. At the participant level, the model parameters were constrained by group-level hyperparameters. We assumed flat distributions for each parameter at the highest level of the hierarchy (hyperparameters).

We used the Gibbs sampler and Markov Chain Monte Carlo technique to perform posterior inference on the parameters in our Hierarchical Bayesian models. To ensure convergence, we drew a minimum of 2,000 samples from an initial burn-in sequence, followed by 5,000 new samples using three chains generated from different random number generators with different seeds. We thinned the resulting sample by a factor of 5 to reduce autocorrelation among the final samples for each parameter, resulting in a final set of 3,000 samples. Gelman-Rubin tests confirmed the convergence of all latent variables in our models, with a statistic near 1 indicating convergence to the target posterior distribution. Due to computational constraints in the TMS-EEG experiment, we estimated individual alpha parameters using 20% of the data (the fifth run of each sub-block). We used this individual alpha parameter to estimate the remaining parameters using the complete data series.

### Anatomical data

All participants in experiments 2 and 3 underwent a 3D anatomical MPRAGE T1-weighted (repetition time [TR]/ echo time [TE]=2530/2.19 ms, inversion time [TI]=1100 ms, flip angle=7°; 1x1x1 mm3 voxels) and T2-weighted (TR/TE=3200/412 ms, flip angle=120°; echo train length [ETL]=258; 1x1x1 mm3 voxels) Magnetic Resonance Imaging scan on a 3T Siemens Skyra scanner (Siemens AG, Erlangen, Germany) with a gradient of 45 mT/m and a maximum slew rate of 200 mT/m/s. The anatomical volume consisted of 160 sagittal slices of an isotropic voxel size (1x1x1 mm), covering the entire brain. The T1-weighted / T2-weighted corrected anatomical MRI was used to extract the scalp and cortical surfaces using a pipeline from the Human Connectome Project, detailed elsewhere^85^. This process yielded a surface triangulation for each envelope,^86^ resulting in individual high-resolution cortical surfaces with approximately 300,000 vertices per surface. These surfaces were then down-sampled to approximately 5,000 vertices. In addition, a five-layer segmentation based on T1-weighted / T2-weighted corrected and T2-weighted images were performed using the algorithm implemented by the SimNIBS tool and SMP12.

### Functional MRI Data

During the fMRI experiment, functional images were acquired using a weighted echo-planar T2* sequence (TR/TE=2390/35 ms, flip angle=90°, 3 × 3 × 3 mm voxels) while participants performed the MSIT task. The acquired volumes of each participant were then coregistered to the 2-mm standard imaging using the nonlinear algorithm, FNIRT, implemented in FSL.^87^ To analyze the imaging data, a model was used that isolated the activity associated with the expectancy of conflict using three regressors of interest: conflict stimuli, non-conflict stimuli, and conflict expectancy (Fig. 2). Second-level activation maps were calculated using a mixed-effects model in FSL, which evaluated two contrasts: conflict higher than no conflict and expectancy of conflict stimuli (Exp=Q, Fig. 2). Cluster correction was applied with a cluster threshold detection (CTD) of z>3.1 and a significance level of p<0.05 using FLAME1.

### TMS-EEG

The stimulation coordinates were individually set to each subject’s brain space in both TMS-EEG sessions. One session was conducted according to the group peak in the SFG for Expectancy regressors, while the other was based on the IFJ for Conflict regressors (Fig. 2). A neuronavigation system was used to identify individual stimulation points (individual structural MRI scans, native space) in the nearest gray matter areas to the no-linear inverse co-registration of the individual anatomy (FSL algorithm, FNIRST) and restricted to be in A6/44d_L (SFG) and IFJ_L (IFJ) of Braintomme atlas segmentation.^43^ TMS coil positioning and orientation regarding the brain x, y, and z axes (that is, yaw, pitch, and roll, respectively) were optimized so that the electric field impacted perpendicular to the target region, maximizing the induced current strength.^65,88^ The stimulation intensity was set to the maximum tolerance level for each subject in this area, ranging from 80% to 120% of their motor threshold previously measured. A 70-mm double coil (PMD70) was used for TMS pulses, which were delivered in three conditions: the theta condition (10 rhythmic pulses every 200 ms), the no-theta condition (10 arrhythmic pulses within 2 seconds), and the sham condition. Sham stimulation was performed in the same area but with the coil tilted, so participants did not receive actual stimulation. EEG recordings were obtained throughout the task in both sessions. We used TMS-compatible EEG equipment (BrainAmp 64 DC, BrainProducts, http://www.brainproducts.com/). EEG was continuously acquired from 64 channels (plus an acquisition reference (FCz) and a ground). TMS-compatible sintered Ag/AgCl-pin electrodes were used. The signal was band-pass filtered at DC to 1000 Hz and digitized at a sampling rate of 5000 Hz. Skin/electrode impedance was maintained below 5kΩ. Electrode impedances were re-tested during pauses to ensure stable values throughout the experiment. The positions of the EEG electrodes were estimated using the neuronavigation system used for the TMS.

### TMS electrophysiological analysis

For both stimulation sites, EEG power was modeled in two-time windows: one between the last TMS pulse, the first Go stimulus, and the other around Go stimuli. These analyses were performed using Matlab software’s LAN package (https://github.com/neurocics/LAN_current) (https://matlab.mathworks.com/).

In the analysis between the last TMS pulse and the first Go stimuli, preprocessing was performed in multiple steps. We first detected the slow decay component of the TMS artifact. To this end, we segmented 3-second windows containing TMS pulses, automatically detected a period starting 10 ms pre to 20 ms post to the respective TMS peak, and removed this from the signal. We applied an Independent Component Analysis (ICA) to this signal using the Runica algorithm provided by the EEGLAB toolbox (https://sccn.ucsd.edu/eeglab/). Thus, we looked for a stereotype component with local bipolar distribution over the TMS site pulse and other psychological signals, such as blinking. In the second step, we segmented the raw signal in the time widows of analysis. We removed the segment between −10 to 30 ms around each TMS stimulation. We replaced it with an inverse-distance weighted interpolation [Y = sum(X/D^3)/sum(1/D^3)] plus a Gaussian noise with the standard deviation extracted to a reference period set to be 55 to −15 ms before the respective first TMS pulse and 0 of the mean. This procedure effectively removed the direct (non-physiological) and other TMS artifacts (e.g., TMS-locked artifacts at electrodes directly in contact with the TMS coil) without introducing discontinuities, essential for the later time-frequency analysis.^62,65^ Following these steps, we down-sampled EEG data to 1000 Hz and used a preprocessing pipeline developed for prior work.^30,31,34,89–92^ The EEG data was then filtered 0.1–45 Hz band-pass and segmented −0.5 to 1.5 seconds around the last TMS pulse. The independent component analysis (ICA) calculated in the first step removed components such as TMS-related artifacts, blinks, and eye movement. Automatic artifact detection was used to remove noisy trials, including voltage threshold (150 μV and three std. dev.) and FFT-amplitude threshold (power spectrum greater than two std. dev. for more than 10% of the 0.5-to-40-Hz spectrum). A final manual review of all trials was performed to check for remaining noise trials, and mastoid electrodes were removed. A Morlet-type frequency analysis (5-cycle width) was performed, and a model with two dummy regressors, Theta TMS and No-Theta TMS, was fitted for each participant.

In the analysis around Go stimuli, data from each participant were preprocessed by decreasing the sampling rate to 1000. Go stimuli were segmented, noisy trials were removed by automatic artifact detection, and ICA was performed to remove blinks and eye movement. Mastoid electrodes were removed, and a manual review of all trials was carried out. The noisy channels were interpolated, and Morlet-type frequency analysis was performed. A model with seven regressors was fitted for each participant: Go, preceding Nogo error, preceding Go error, Expectancy, Expectancy*TMS interaction, Expectancy*TMS_theta_ interaction, TMS, TMS_theta_. Second-level analysis was performed in both time windows by incorporating the models of each participant to model the power according to EEG and analyze brain electrical activity according to areas, conditions, times, and frequencies.

### Cluster-based Permutation Test

We employed a cluster-based permutation test to address multiple comparisons in our time-frequency analysis.^93^ Significant regions were identified by clustering neighboring sites showing the same effect (with a significance level of p < 0.05 in the statistical tests applied to either the time-frequency chart or the sources, such as the Wilcoxon test). Cluster-level statistics were computed as the sum of statistics from all sites within the cluster. We evaluated the significance of these clusters by comparing them to the largest cluster-level statistics in a permutation distribution. This distribution was generated by randomly permuting the original data. Specifically, we created null models for each participant, maintaining the original model structure while permuting the tested regressor. Following each permutation, we conducted the initial statistical test (e.g., Wilcoxon) and recorded the cluster-level statistics of the largest cluster from the permutation distribution. After 5,000 permutations, we estimated the cluster-level significance for each observed cluster as the proportion of values in the permutation distribution surpassing the cluster-level statistics of the corresponding cluster

### EEG Source Estimation

We estimated the neural current density time series at each brain location using a minimum norm estimate inverse solution LORETA algorithm with unconstrained dipole orientations. This estimation was performed individually for each trial, condition, and subject, employing Brainstorm software^94^. We used a tessellated cortical mesh to model the brain based on each individual’s anatomy. This mesh was employed to estimate the distribution of current sources, with approximately 3×8000 sources positioned on the segmented cortical surface (three orthogonal sources at each spatial location). We employed a five-layer continuous Galerkin finite element conductivity model (FEM), as implemented in DUneuro software^95^, along with a physical forward model. We multiplied the recorded raw EEG time series from the electrodes by the inverse operator to obtain cortical activity estimates at the cortical sources. This operation yielded the estimated source current at the cortical surface as a function of time. Importantly, this transformation is linear and does not alter the frequencies of the underlying sources, enabling us to conduct time-frequency analyses directly in the source space. Within this source space, we simplified each vertex’s dipole to one by selecting the component with the highest variance, employing the PCA algorithm. Subsequently, we performed frequency decomposition using the Wavelets transform. To ensure the reliability of our results, we only present source estimations if statistically significant differences are observed at both the electrode and source levels, thereby accounting for multiple comparison corrections.

## Data Availability

The complete minimal data set underlying the results used in our study will be available in the public repository of the Openneuro once accepted for publication― (doi: xxxx). The additional toolbox and codes used in the analysis are available on our lab github site at https://github.com/neurocics/ and the OSF website xxxx.

## Supplementary Materials

**Supplementary Fig. 1.**
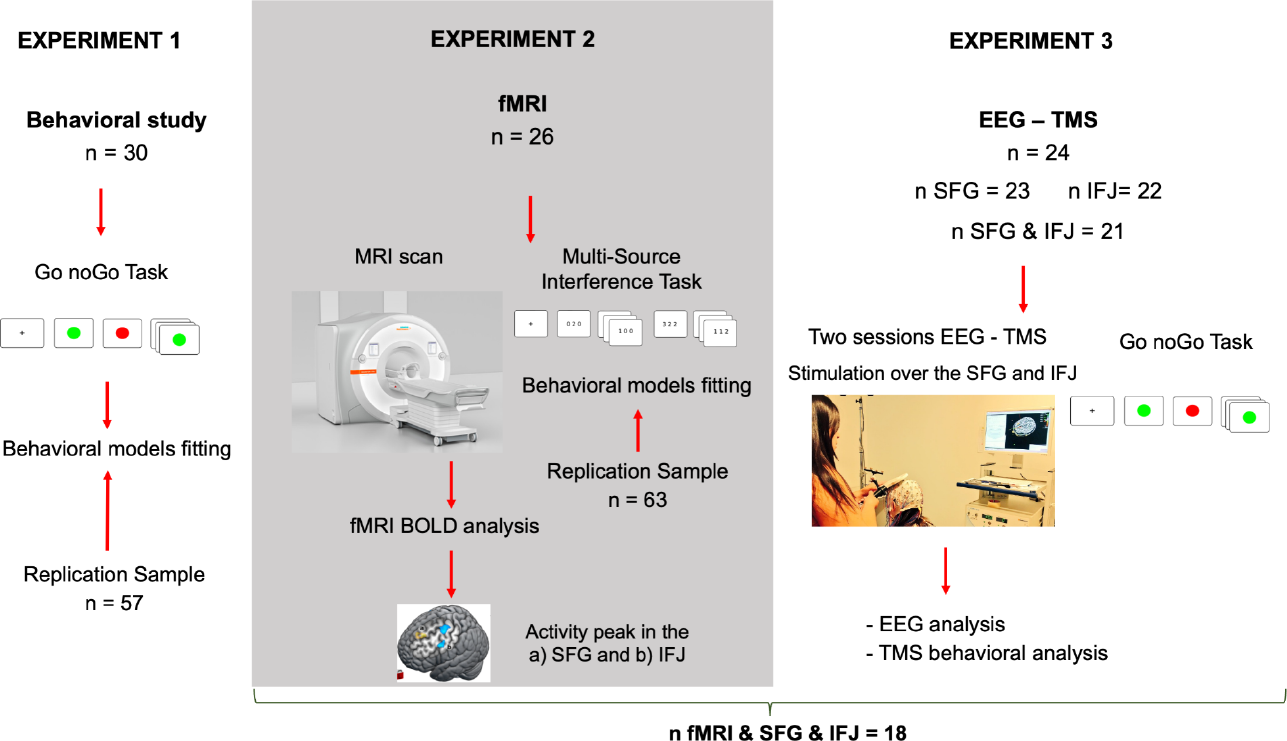
Experimental design. Three experiments were carried out. Experiment 1: Behavioral study (n=30) with GNG task. Experiment 2: fMRI session with MSIT task (n=26). Experiment 3: Two randomized EEG-TMS sessions, one in the SFG and other in the IFJ with GNG task (n=24). GNG: Go Nogo; fMRI: Functional Magnetic Resonance Imaging; SFG: Superior Frontal Gyrus; IFJ: Inferior Frontal Junction; EEG: Electroencephalography, TMS: Transcranial Magnetic Stimulation.

**Supplementary Fig. 2.**
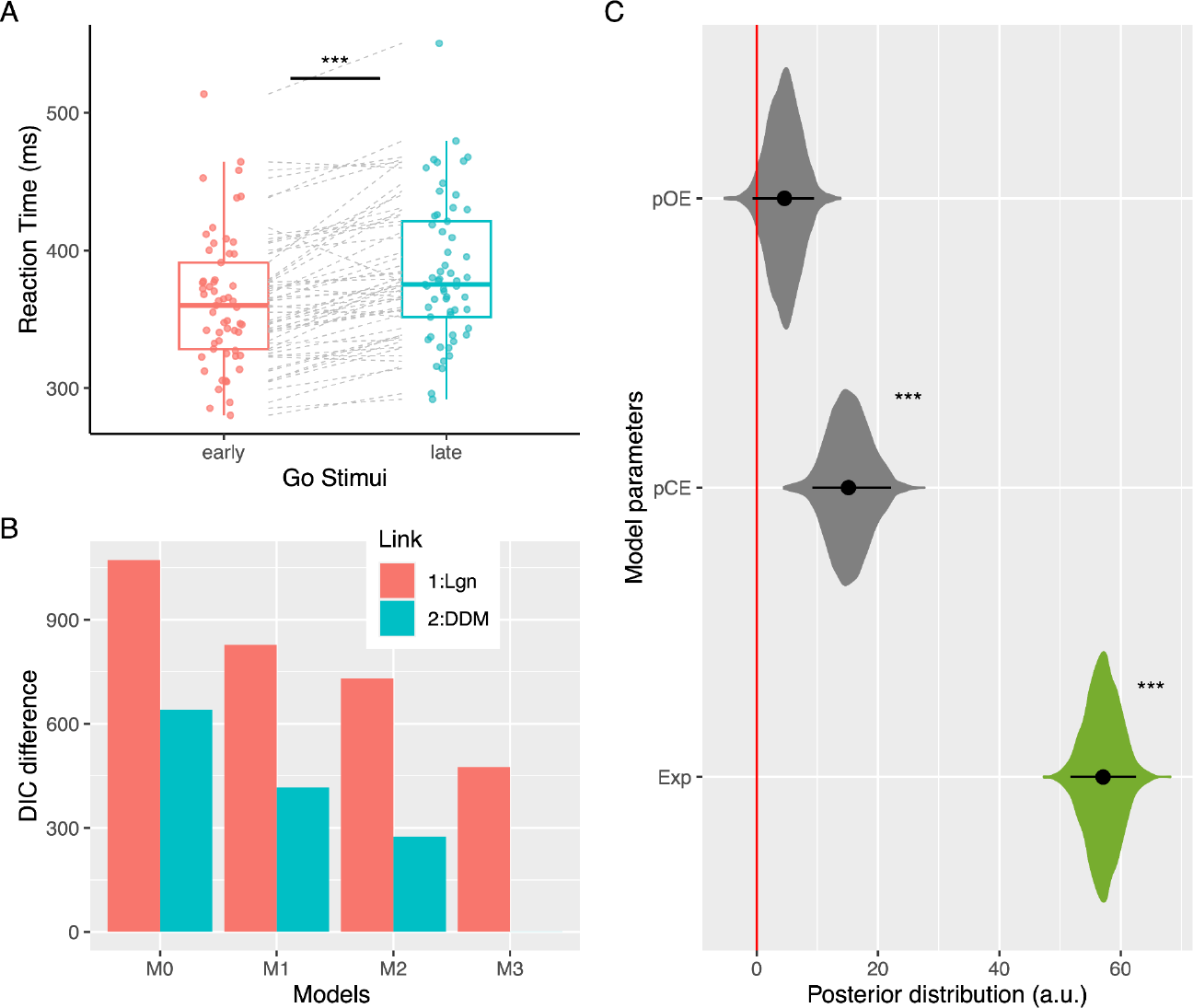
GNG replication sample. Behavioral analysis of GNG Task replication sample. A) Reaction time comparison between the first Go trials (early) and the last Go trial (late) of a sequence. B) Models comparison. C) Posterior distribution of model parameters. DIC: Deviance Information Criteria. Exp: Conflict Expectation. pCE: Previous Commission Error. pOE: Previous Omission Error. RT: Reaction Time. See also Supplementary Table 1.

**Supplementary Fig. 3.**
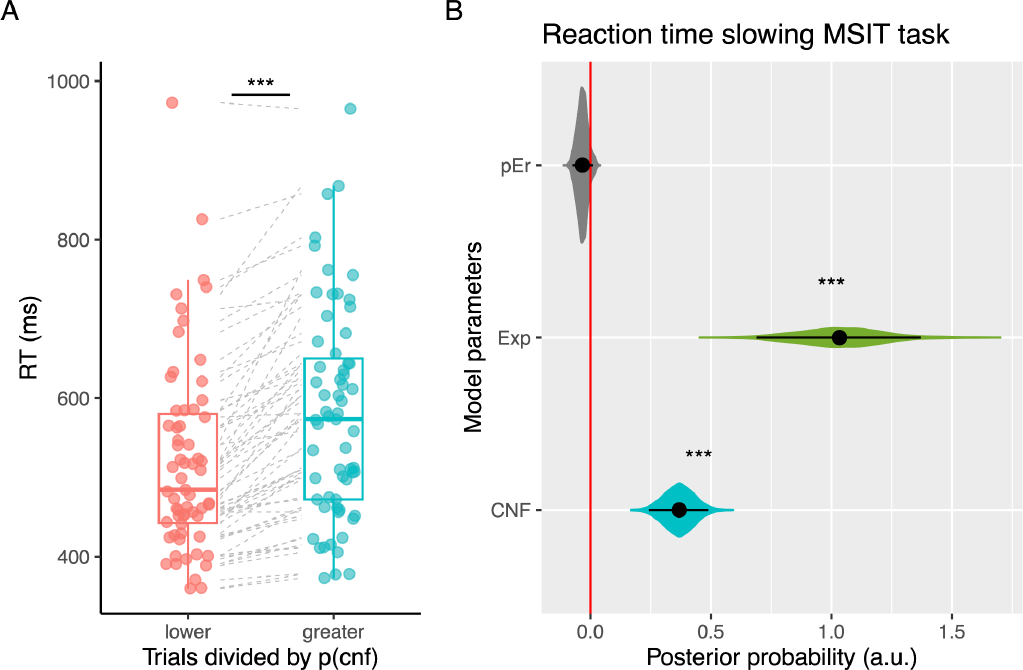
MSIT replication sample. **A**. Comparison of reaction times between trials with predicted lower and higher conflict expectations per sequence. Each point represents the mean of a subject. **B**. Posterior distribution of model parameters. pEr: Error in the prior trial, Exp: Conflict Expectation, CNF: Conflict trial, p(cnf): Predicted probability of conflict, RT: Reaction Time.

